# ROIMAPer: An Open-Source Framework for Rapid and Accurate Atlas-Based Registration of Individual Brain Images in FIJI

**DOI:** 10.64898/2026.03.16.712226

**Authors:** Julian N. Rodefeld, Annie V. Ciernia

## Abstract

The brain’s remarkable complexity and cellular heterogeneity necessitate precise anatomical annotation to ensure that imaging-based analyses accurately resolve region-specific features. Few computational tools currently exist that allow for the accurate and rapid registration of single brain images to standard brain atlases. To address this limitation, we developed ROIMAPer, a novel FIJI plugin for rapid registration of individual brain slices. ROIMAPer includes eight atlases spanning mouse, rat, and human brain anatomy across multiple developmental stages, making it broadly applicable across diverse experimental contexts. It allows for linear and affine scaling of the reference atlas to the experimental image and is optimized for serial processing of large quantities of images. We demonstrated the accuracy of ROIMAPer through quantification of *in situ* hybridization data from the Allen Gene Expression Atlas of seven marker genes across major brain regions and of four marker genes across hippocampal subfields. Quantification of marker genes within their assigned brain regions closely matched the ground truth across all major regions. At a finer resolution, marker-gene quantification within hippocampal subregions aligned with the experimental data, although discrepancies with the ground truth were observed for *Mcu*. Overall, ROIMAPer provides broad utility for open-source brain image analysis from multiple species.

**Significance Statement:** We present an open-source, user-friendly, and accessible tool for registration of individual brain slices to anatomical reference atlases, compatible with the image analysis platform FIJI. The field lacks tools that offer a span of cross-species atlases, FIJI-compatibility, intuitive linear scaling methods, and low user-input without requiring high computational skill. Our tool minimizes user-involvement, allows for processing of larger datasets through more effective resource management, and speeds-up previously tedious processing steps.

## Introduction

Neurological assessment commonly depends on imaging techniques such as MRI, immunohistochemistry, or *in situ* hybridization. To interpret these data in a region-specific manner and to capture the brain’s pronounced anatomical heterogeneity, images must be annotated with the relevant anatomical regions. This process, referred to as brain registration, is performed by aligning an experimental image to a standardized reference brain atlas. Reference atlases are typically constructed by aggregating morphological information across many individuals to generate a representative anatomical template (Toga and Thompson, 2001). In widely used resources, this may involve MRI complemented by high-resolution microscopy (Hawrylycz et al., 2014) or serial two-photon tomography (Allen Institute for Brain Science, 2017). The resulting template is then annotated with specific brain regions based on anatomical features or region-specific gene expression. Accurate registration to such atlases enables quantitative, reproducible comparisons across experiments and subjects.

The major challenge during the alignment of an experimental brain image to the reference atlas is the correct position of an image within the atlas and compensating for tissue deformation. Key cues for atlas positioning include the recorded section order (e.g., section number noted during cutting), the overall shape of the tissue section—which changes systematically along the anterior–posterior axis—and identifiable anatomical landmarks such as major white-matter tracts. In practice, variability in sectioning, mounting, and staining can introduce distortions, so experimental sections may deviate from the idealized atlas geometry and require local adjustment to achieve accurate alignment.

Numerous registration tools approach these positioning and deformation steps through a range of strategies. Many of these were developed to register serial sections that compose large portions of the brain (Puchades et al., 2019, 2025; Carey et al., 2023; Soronow et al., 2025). However, numerous experimental workflows, such as immunofluorescence studies, often involve processing individual slices drawn from different brains, where rapid and flexible single-section registration is required. Several tools address this problem, but require advanced programming skills, large degrees of manual input, are difficult to install or use, or not compatible with the open-source image analysis platform FIJI/ImageJ (Schindelin et al., 2012). For example, Brain-Ways (Kantor and Bartal, 2023) can provide an initial atlas position estimation based on a machine learning model and allows for manual adjustment via non-rigid registration using Elastix (Klein et al., 2010; Shamonin et al., 2014). But its linear scaling method is difficult to master, and its output is not compatible with FIJI/ImageJ. AMBIA (Sadeghi et al., 2022, 2023) only includes an atlas for the adult mouse and has a complicated installation process. The FASTMAP plugin (Terstege et al., 2022) for FIJI/ImageJ was the direct inspiration for the development of our tool. It aims for highly flexible image registration by relying on user-specified atlases dependent on the experimental needs, however, it requires extensive manual user-input during the adjustment of the registration and saving processes.

We aimed to develop a new tool that is easy to install, allows processing with minimal manual input, and is FIJI/ImageJ compatible. Here, we present ROIMAPer, a FIJI-plugin that enables rapid brain registration of individual brain slices, supports efficient manual refinement via affine Delauney triangulation, and includes eight widely used atlases spanning different species and developmental stages.

## Materials and Methods

### Brain Atlas Generation

The rat atlas was processed in FIJI while the mouse and human atlases were processed using version 2.4.0 of SimpleITK (Lowekamp et al., 2013; Yaniv et al., 2018) in python (version 3.11.5) (“Python Language Reference, version 3.11.5,” n.d.). All atlases were resliced in coronal and sagittal direction, with slicing intervals emulating common 2-dimensional reference atlases (Table 1). To prevent inaccurate 32-bit float conversion upon import of the atlases in ImageJ/FIJI during usage of the plugin, the modulo 100 000 of the pixel values of all mouse atlases was taken. For the human atlas the modulo 1 000 000 was taken, to prevent duplicates indices. Coronal versions of the atlases were bisected vertically to supply an atlas for mapping of single hemispheres.

**Table 1:** Altas slicing specifications. The slicing range describes the first and last image of the z-stack that was used to create the final atlas in FIJI. The slicing interval specifies how frequently slices from the z-stack were sampled for the final output. The P56 mouse brain is represented both through adult and developmental nomenclature – allowing for cross-comparisons between the two systems.

|  | Coronal |  | Sagittal |  |
| --- | --- | --- | --- | --- |
| Atlas | Slicing range | Slicing interval | Slicing range | Slicing interval |
| Adult mouse brain | 3-1320 | 10 | 1-570 | 20 |
| P56 mouse brain | 3-1320 | 10 | 1-570 | 20 |
| Adult human brain | 54-416 | 4 | 52-197 | 5 |
| Adult rat brain | 2-1024 | 10 | 2-512 | 10 |
| E11.5 mouse brain | 0-326 | 3 | 0-71 | 1 |
| E13.5 mouse brain | 0-391 | 3 | 0-87 | 2 |
| E15.5 mouse brain | 0-464 | 4 | 0-122 | 2 |
| E18.5 mouse brain | 0-581 | 4 | 0-145 | 2 |
| P4 mouse brain | 0-724 | 5 | 0-193 | 3 |
| P14 mouse brain | 0-794 | 6 | 0-195 | 3 |
| P28 mouse brain | 0-863 | 6 | 0-200 | 3 |
| P56 mouse brain (developmental brain nomenclature) | 0-528 | 4 | 0-228 | 4 |

All atlases were obtained together with their respective brain region ontology files. Within these, the intensity values in the image, corresponding to the local brain region index, are mapped to the names and acronyms of the respective brain regions. In case of nested regions, hierarchy information in the form of the index of the parent region is included as well. Since this list is used to look up the index of requested brain regions, the modulo conversion that was performed on the images themselves was repeated in these lists, maintaining consistency of the indices. An overview of every atlas in every available orientation was created, using false colouring of each possible slice.

The first time a brain region is requested in ROIMAPer, the corresponding index is identified in the ontology file. If this brain region is composed of different subregions, their indices are gathered in an array. The atlas image is then thresholded for each index in that array, forming a mask for every subregion and the parent region. All the resulting binary masks are combined to form a 3-dimensional mask of the full brain region. Finally, each 2-dimensional slice of the mask is transformed into an ImageJ “region of interest” (ROI).

To establish a scaling reference, a bounding box is defined as a rectangular selection encompassing the full extent of the brain and saved together with the brain region ROI. A record is kept of which brain regions have been created in each atlas, preventing the mask from being regenerated on subsequent requests.

### Scaling and Rotation of Experimental Images

All experimental images that are to be processed are indexed at the beginning of the ROIMAPer registration. If blinding is desired, the order of the images is randomized and the name of every opened image is obscured. Only one slice and one registration channel of the experimental image is opened to increase performance and decrease memory strain. A bounding box around the experimental brain in this registration channel is either created by the user or using the ImageJ “Fit Rectangle” algorithm. The ROI.zip files of the atlas brain regions of interest generated as described above are opened and are first linearly scaled before they are moved and rotated to proportionally fit inside the experimental bounding box. The scaled ROIs can be flipped in either x or y direction, as well as rotated by 90°, if the bounding box was created in a different orientation relative to the atlas. After one image has been processed this way, the mapped image is either saved immediately or only the ROI.zip file is saved in a temporary folder and the actual saving of the images is performed after all images have been registered, minimizing wait times between images.

### Affine Mesh Transform

To allow for easy manual correction of the linear scaling, we implemented affine deformation (Sotiras et al., 2013) based on Delauney triangulation, which has been used in brain registration before (Puchades et al., 2025). A user-specified collection of points and the four points outside the corners of the brain bounding box are combined to form the vertices of the transformation mesh. For every point *P* in the border of every brain region ROI, the three closest transformation mesh vertices *A, B, C* are found, that both include the point *P* in their formed triangle and include no other vertex in their circumcircle, resulting in the enclosing Delauney triangle.

From these three vertices, their barycentric weights are calculated according to:

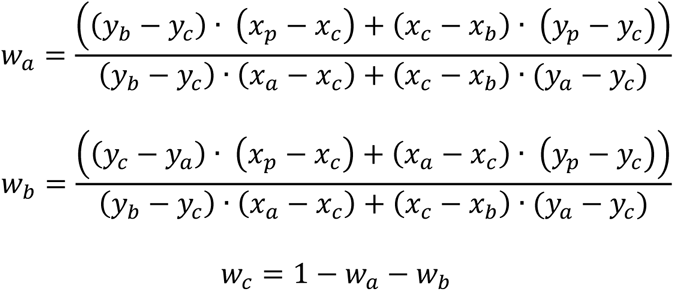

After translation of *A, B, C* to *A’, B’, C’* by the user, the coordinates of *P’*, the new point of the ROI border, are calculated as:

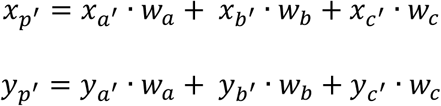

For composite ROIs, the individual components are separated before the transformation, processed independently, and then recombined afterwards.

### Quantification of expression energy in ISH images

*In situ* hybridization (ISH) images and images showing their summarized expression energy for eleven genes were batch downloaded from the Allen Gene Expression Atlas (AGEA) (Lein et al., 2007) (Table 2). Brain regions were registered using the ROIMAPer on the ISH brightfield images and were transferred to the red channel of the ISH expression energy summary. The mean intensity of the red channel was quantified for each region in ImageJ. The ground truth of the expression energy was retrieved from the Allen Brain Atlas API. The mean expression intensity per gene was averaged per region, resulting in one datapoint for every brain region.

**Table 2:**
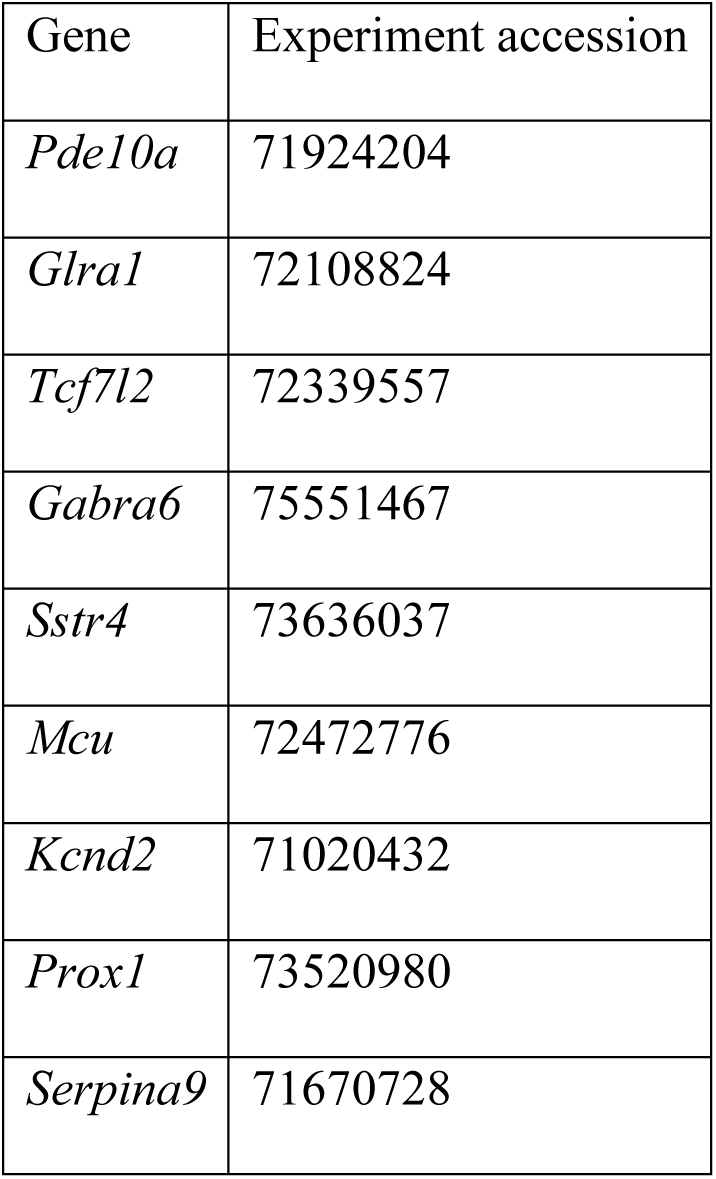
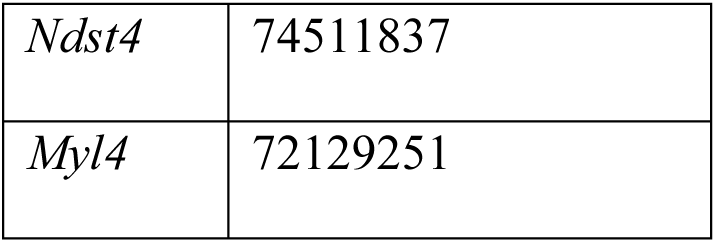
Accession numbers for ISH records available from https://mouse.brain-map.org/experiment/show/[accession].

### Statistics, Data Management and Figures

To test for similarity between the ground truth and the ROIMAPer quantification, common statistical tests like t-test or ANOVA are unfit, since they only test for differences. Instead, a paired two one-sided test of equivalence (Schuirmann, 1987) with TOSTER (version 0.8.6) (Lakens, 2017; Caldwell, 2022) was performed between the ISH expression signal from the Allen Institute and ROIMAPer quantification for each gene, across either the 12 major brain regions or four hippocampal subregions.

We converted the expression energy into relative expression energy by dividing through the total expression in every brain and used an equivalence bound of θ = 5 % according to (Limentani et al., 2005). The p-value was corrected for multiple testing across different genes with the ℓ-correction (Lauzon and Caffo, 2009).

Data organization was performed and graphs were generated in R (version 4.5.2) (R Core Team, 2025), using dplyr (version 1.1.4) (Wickham et al., 2023) and ggplot2 (version 4.0.1) (Wickham, 2016). Final figures were created in Inkscape v1.4.2.

### Code Availability

The code/software described in the paper is freely available online at https://github.com/ciernialab/ROIMAPer or through the FIJI Updater on the update site https://sites.imagej.net/ROIMAPer/. The code is available as Extended Data.

## Results

### Brain Atlas Preparation

When performing imaging-based methods like immunofluorescence microscopy, the registration of brain regions across images is vital for downstream distinction of region-specific effects. The following describes the development and assessment of ROIMAPer, a FIJI plugin for easy, rapid, and accurate single-slice brain registration across multiple species.

ROIMAPer requires a brain atlas for registration. To allow for flexible mapping to mouse, rat and human brain images, we collected atlases for the adult mouse (Allen Institute for Brain Science, 2022), postnatal day 56 (P56) mouse (Allen Institute for Brain Science, 2016), and the developing mouse brain (Young et al., 2021) at embryonic day 11.5, 13.5, 15.5, and 18.5, and postnatal day 4, 14, 28, and 56. The Waxholm space atlas of the Sprague Dawley rat (Papp et al., 2014; Kleven et al., 2023) was used for adult rats and the adult human brain from (Ding et al., 2020). The atlas files were prepared to be compatible with ImageJ, and a selection of slices across both the coronal and sagittal axis were created.

### Process of ROIMAPer Registration of Brain Images

The main goal of ROIMAPer is fast registration of individual brain images in FIJI. To maximize processing speed and minimize memory performance, brain region registration is performed on only one channel of the experimental image. After specifying the experimental images to register, the user selects the desired brain regions (Figure 1A), estimates the appropriate position in the atlas from an overview (Figure 1B), and either manually aligns a bounding box around the brain or allows for the automatic alignment of this bounding box (Figure 1C). ImageJ ROI objects are created from the atlas files and are linearly scaled, moved, and rotated to the bounding box. Manual refinement of the brain regions can be performed through mesh-based Delauney triangulation (Figure 1D). This prompts the user to first plant landmark points on the image in places where adjustment is necessary, and to then move them to the new, appropriate positions, resulting in dynamic adjustment of the brain regions.

**Figure 1:**
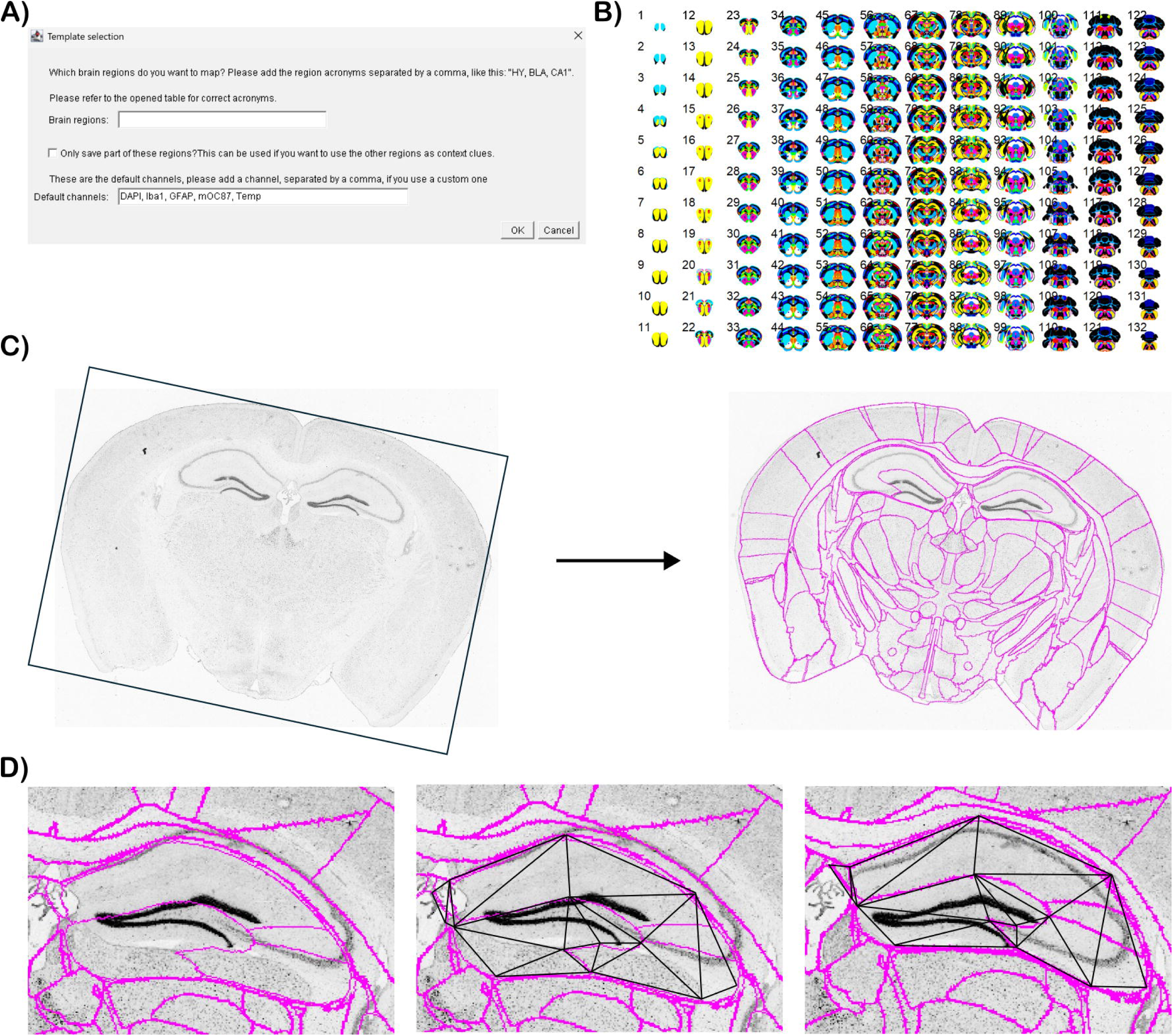
Workflow of the ROIMAPer FIJI plugin. A) Example of the FIJI user interface, asking for region and channel specification. B) Overview atlas of the adult mouse brain. C) Initial ROI registration with linear scaling to the brain bounding box. H) Delauney triangulation for brain region adjustment (hippocampus) after linear scaling, based on user-specified points. Brain images were obtained from https://brain-map.org.

To reduce bias during processing, image names can be blinded, and image order can be randomized. The saving of image data is often time-intensive, so saving can be performed after all images are processed. This user-friendly plugin enables registration of multiple brain regions within a single image and rapid processing of multiple images.

### Accurate Brain Registration to a Reference Atlas with ROIMAPer

We assessed the accuracy of ROIMAPer by working with *in situ* hybridization images (ISH) from the Allen Gene Expression Atlas (AGEA) (Lein et al., 2007). The expression of seven genes was analysed across a total of 276 images and averaged across twelve major brain regions: Isocortex, olfactory bulb (OLF), hippocampus (HPF), cortical subplate (CTXsp), striatum (STR), pallidum (PAL), thalamus (TH), hypothalamus (HY), midbrain (MB), pons (P), medulla (MY), and cerebellum (CB). The expression values were then compared to the original quantification (Lein et al., 2007) as a ground truth.

The similarity between quantification with ROIMAPer and the ground truth was assessed using a two one-sided test of equivalence (TOST), which tested whether two sets of data were significantly similar within certain bounds (Schuirmann, 1987). The sample-size was defined as the compared number of regions. A p-value lower than the significance level of α = 0.5 showed that the datasets were equivalent.

We examined a series of marker genes with unique brain region specific expression (Figure 2). For *Myl4* (Figure 2A) the two datasets were equivalent across all brain regions (t-test: t(11) = - 2.09e-16, p = 1; equivalence test: t(11) = 2.257, p = 0.023). For *Ndst4* (Figure 2B) the two datasets were tested as equivalent (t-test: t(11) = -4.12e-16, p = 1; equivalence test: t(11) = 5.084, p < 0.001). *Pde10a* (Figure 2C), a marker gene for the striatum (Seeger et al., 2003; Lakics et al., 2010), was equivalent between the ground truth and ROIMAPer (t-test: t(11) = - 1.15e-15, p = 1; equivalence test: t(11) = 6.231, p < 0.001). The serine protease inhibitor 9a (*Serpina9*) (Figure 2D) was tested as not equivalent between our quantification and the ground truth (t-test: t(11) = 3.67e-16, p = 1; equivalence test: t(11) = -1.63, p = 0.066). *Tcf7l2* (Figure 2E), a marker for the thalamus (Nagalski et al., 2013, 2016) and the brain stem/midbrain (Lee et al., 2009), was equivalent between the ROIMAPer quantification and the ground truth (t-test: t(11) = 1.39e-17, p = 1; equivalence test: t(11) = -4.74, p < 0.001). *Glra1* (Figure 2F) is highly expressed in the midbrain (MB), pons (P), and medulla (MY) (Malosio et al., 1991; Callister and Graham, 2010). The quantification of the expression of *Glra1* between the two datasets was equivalent (t-test: t(11) = -1.22e-15, p = 1; equivalence test: t(11) = 7.949, p < 0.001). Finally, the expression of *Gabra6* (Figure 2G), a cerebellar marker gene (Wisden et al., 1992), was equivalently quantified between both datasets (t-test: t(11) = -2.01e-16, p = 1; equivalence test: t(11) = 2.485, p = 0.015).

**Figure 2:**
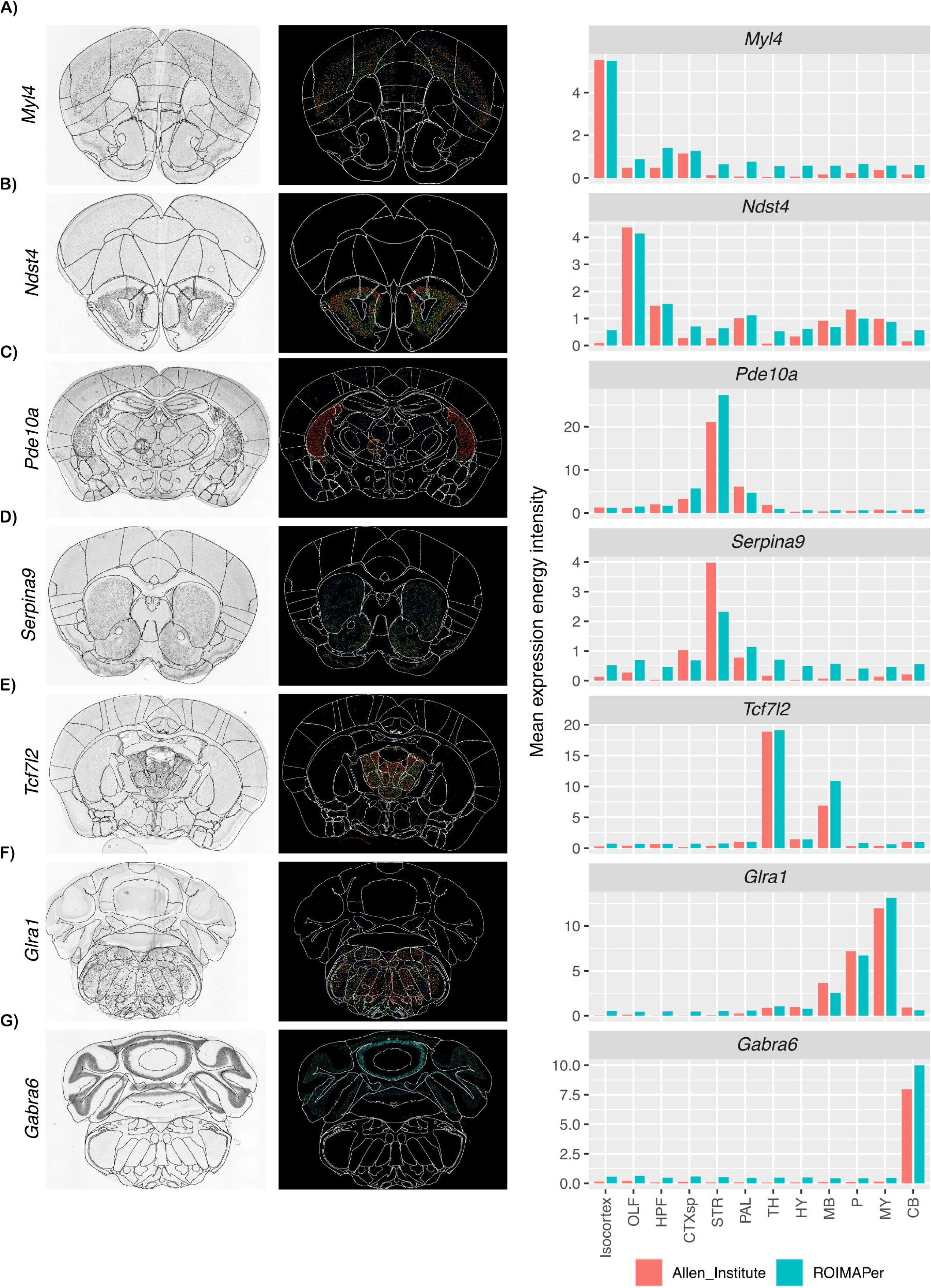
ROIMAPer captures the brain region specific Allen Institute ISH data across large regions. Comparison of the original ISH signal across the brain regions with signal after brain region registration using the ROIMAPer. Greyscale of a representative brain slice with ROIMAPer regions drawn in black are shown alongside RGB colorations of the ISH expression energy of that same representative brain slice with the ROIMAPer regions in white. Each graph shows the quantification of the expression energy, comparing the brain regions registered using the ROIMAPer (turquoise) and the original region signal of the Allen institute (red). A) *Myl4*. B) *Ndst4*. C) *Pde10a*. D) *Serpina9*. E) *Tcf7l2*. F) *Glra1*. G) *Gabra6*.

To gain insight at a higher level of detail, subregions of the hippocampal formation were investigated in a similar manner across 66 images. Four marker genes for the major subregions of the hippocampus were chosen: The somatostatin receptor 4 (*Sstr4*) (Figure 3A) which is most specific to the CA1 field of Ammon’s horn (Meyer, 2014), with some claims to expression in all of Ammon’s horn (Gastambide et al., 2010). *Mcu* (Figure 3B) is a marker gene for the CA2 field (Farris et al., 2019) and *Kcnd2* (Figure 3C) has been identified as a specific marker for the CA3 field (Bugaj et al., 2024). Finally, *Prox1* (Figure 3D) is a marker specific to the Dentate Gyrus (DG) (Machado et al., 2022). None of the quantifications were equivalent to the ground truth, but the test was underpowered since there were only four points of comparison each, limiting its interpretability: *Sstr4* (t-test: t(3) = 1.373, p = 0.26; equivalence test: t(3) = 1.27, p = 0.85), *Mcu* (t-test: t(3) = 1.876, p = 0.16; equivalence test: t(3) = 1.77, p = 0.91), *Kcnd2* (t-test: t(3) = 1.83, p = 0.16; equivalence test: t(3) = 1.72, p = 0.91), and *Prox1* (t-test: t(3) = - 0.456, p = 0.68; equivalence test: t(3) = -0.31, p = 0.61). The trends between the two datasets were similar in all genes, except for *Mcu*, where the Allen Institute ground truth disagreed with its CA2 marker gene status.

**Figure 3:**
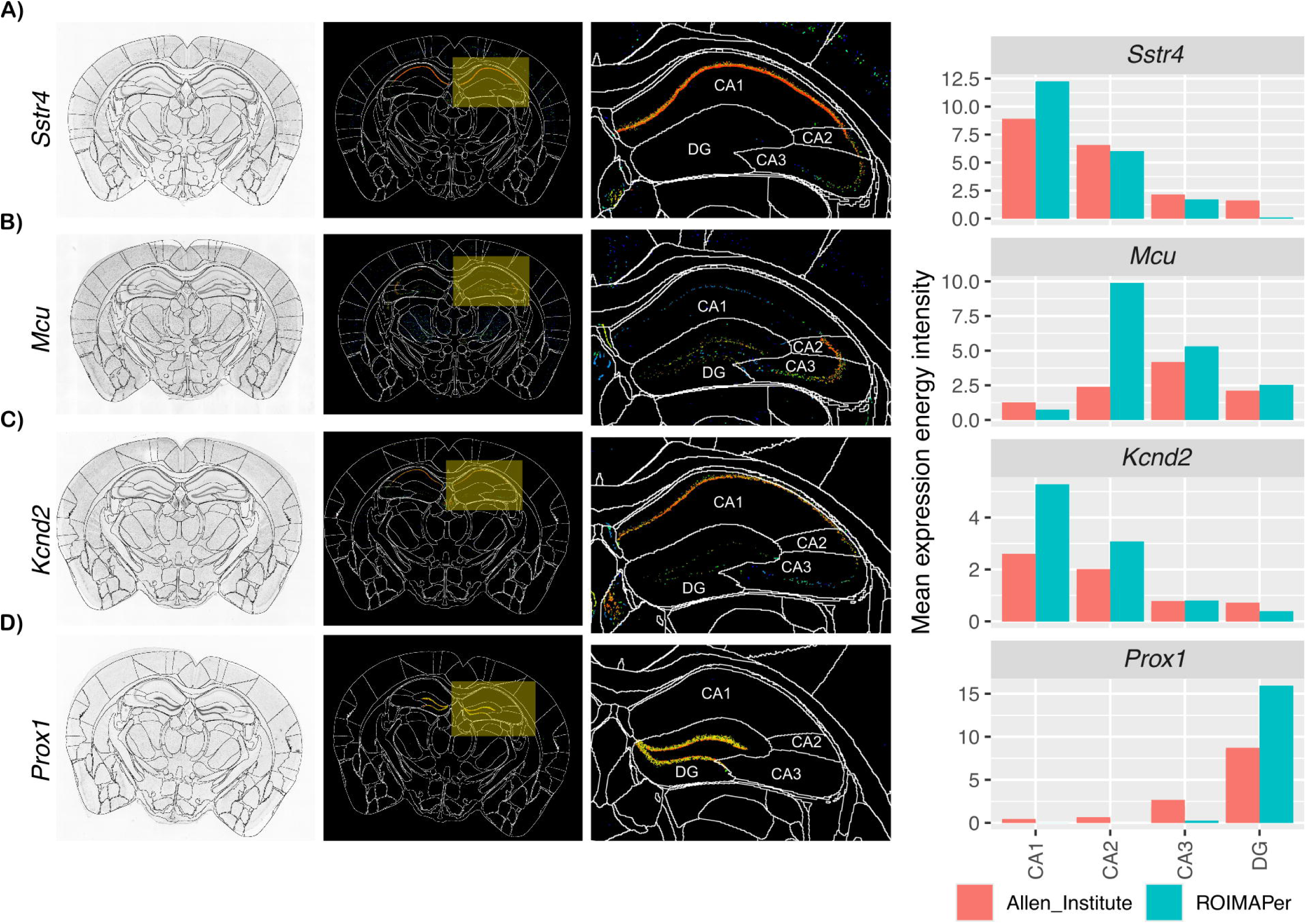
ROIMAPer captures the brain region specific Allen Institute ISH data across hippocampal subfields. Mapping of the hippocampal subfields CA1, CA2, CA3, and DG across ISH images for four marker genes. Shown are greyscale of a representative brain slice with ROIMAPer regions drawn in black, RGB representations of the ISH expression energy zoomed in on the hippocampal formation and the quantification of the expression energy, comparing the ROIMAPer and the quantification of the Allen institute. A) *Sstr4.* B) *Mcu.* C) *Kcnd2.* D) *Prox1*.

This qualitative agreement between ROIMAPer quantification and the Allen Institute ISH quantification on a high level of detail, and the quantitative equivalence between larger brain regions demonstrates that ROIMAPer can be used for accurate and comprehensive brain registration.

## Discussion

In this study we have presented ROIMAPer, a FIJI plugin for the rapid registration of individual brain slices in human, rat, and adult and developing mouse. Brain regions are scaled linearly and can be modified with point-based affine transformation. The tool is easily installable through its ImageJ update site, making it available without advanced coding skills. We have shown that ROIMAPer can accurately map major and minor brain regions based on a gene expression ground truth.

### A Comparison Across Tools for Brain Registration

While there are other tools for brain registration, ROIMAPer has several distinct advantages. First, ROIMAPer is ideally suited for single images that are not done in serial. Landmark-based registration algorithms are poorly suited for single-slice registration because an individual slice may not contain the landmarks required for alignment, and no adjacent slices are available to provide additional reference points. Previous tools such as ABBA (Chiaruttini et al., 2025) register full serial section datasets and depend on contextual clues from multiple neighboring slices, making it unsuitable for workflows that lack serial sections. Additional tools that have the same caveat of being designed for full brain / serial slice registration are: brainreg (Niedworok et al., 2016; Claudi et al., 2020; Tyson et al., 2022), bell jar (Soronow et al., 2025) and VisuAlign (Gurdon et al., 2024; Puchades et al., 2025), an algorithm in QuickNii (Puchades et al., 2019).

Second, ROIMAPer allows for easy batch processing of large sets of images without deep computational knowledge. This has advantages over tools like web version of DeepSlice (Carey et al., 2023) that requires repeated file-upload, which is cumbersome for large datasets. The local version requires basic knowledge in Python, resulting in a barrier of entry for non-computational scientists.

Other dedicated tools for individual registration of coronal brain slices AMBIA (Sadeghi et al., 2022, 2023) and BrainWays (Kantor and Bartal, 2023) offer an initial AI estimate of the brain registration. However, in comparison to AMBIA, ROIMAPer offers an easier installation for users unfamiliar with virtual environments, and a wider selection of atlases. Additionally, AMBIA is only applicable in coronal brain sections. In comparison to BrainWays, the ROIMAPer plugin offers compatibility with FIJI region objects and a more intuitive linear scaling using the bounding box. ROIMAPer offers flexibility in sectioning direction and applicable species. The FASTMAP (Terstege et al., 2022) FIJI plugin was developed for the registration of diverse tissue types and stains to user-created atlases. It sequentially opens each image in the experimental dataset and roughly scales the corresponding atlas regions to the size of the experimental brain, after which the alignment can be manually refined. ROIMAPer offers several advantages over FASTMAP. ROIMAPer has more stable scaling of the brain regions and works with images rotated in every direction. It is also faster and less memory intensive, more automated, and includes fully prepared coronal and sagittal atlases for mouse, rat, and human. Affine transformation via Delaunay triangulation enables flexible adjustment of the atlas to accommodate warped or anatomically altered images, whether due to acquisition artifacts or experimental models.

### Validation of ROIMAPer Accuracy

In major brain regions where high marker-gene expression was expected, the intensity of the expression signal closely matched between the ground truth and the ROIMAPer quantification for all genes. *Serpina9* where the datasets were not equivalent was lowly expressed overall, leading to discrepancies in measurements due to the semiquantitative expression scale. Accuracy across fine brain regions was examined in the hippocampal formation. ROIMAPer demonstrated the presence of the marker genes *Sstr4* and *Prox1* in CA1 and DG, respectively, and demonstrated that previous literature regarding *Kcnd2* being a CA3 specific marker gene (Bugaj et al., 2024) did not agree with the available ISH data. Finally, we showed a quantification of the hippocampal *Mcu* expression that better corresponds to the ISH images and its role as a CA2 marker (Farris et al., 2019) than the quantification of the Allen Institute. These findings support the specificity and accuracy of ROIMAPer to register major brain regions within the test images.

### Limitations and Future Directions

Our quantification of the ISH expression energy was limited by trying to recreate it from the semiquantitative RGB summary. We opted to quantify the signal in the red channel, since it increased roughly linearly across the expression scale. Additionally, the expression energy data was supplied in jpeg format, causing potential data loss due to compression. However, even given these limitations, the majority of ROIMAPer intensity values corresponded closely to the reported ISH values.

Future directions for ROIMAPer include the possibility of a tilting angle of the atlas, to correct for an off-centre cutting angle during sectioning. Additionally, expanding the set of available atlases to include a broader range of species and developmental stages would substantially increase ROIMAPer’s utility. This could be done using the BrainGlobeAtlas API (Claudi et al., 2020), which provides standardized access to a wide collection of atlases and streamlines their use across tools. Creating an additional tool for easy construction of user-specific atlases would allow for use of the ROIMAPer in a wider range of contexts.

## Conclusion

This work developed and validated the ROIMAPer FIJI plugin that provides rapid registration of collections of individual brain images to a reference atlas. ROIMAPer accurately identified regions and subregions across the entire brain with high accuracy. This new tool will enable faster and more accurate processing of large quantities of brain image data across species and ages.

Extended Data 1: Code/Software

Included in this software is the full ROIMAPer program and code that was used to quantify its accuracy.

ROIMAPer.zip: Zipped archive of the ROIMAPer code, including: ROIMAPer.ijm, the FIJI/ImageJ plugin. atlases/ the collection of atlas files included in the plugin. ROIMAPerUtilities/ a collection of ontology files, atlas overview files, and scripts that were used to create said overview and atlas files. images/ supporting images for the README.md file. in_situ_hybridization_download.R: script used to batch access the AGEA files. in_situ_hybridization_fusing.ijm: script used to combine the original ISH images with the semiquantitative expression energy summary for the purpose of registration. in_situ_hybridization_measure.ijm: script used to measure the red intensity of the semiquantitative expression energy after registration with ROIMAPer.

ROIMAPer_quantification.R: script used to compare the ROIMAPer quantification with the ground truth of the AGEA.

