## Extended Data 1 for "ROIMAPer: An Open-Source Framework for Rapid and Accurate Atlas-Based Registration of Individual Brain Images in FIJI": in_situ_hybridization_download.R

```
#download allen brain expression data

start_url = "http://api.brain-map.org/api/v2/image_download/"
genes = c("pde10a", "serpina9", "glra1", "myl4", "tcf7l2", "ndst4", "gabra6", "prox1","mcu", "kcnd2", "sstr4")

experiments = setNames(c(71924204,71670728,72108824,72129251,72339557,74511837,75551467,74881547,72472776,71020432,73636037), 
                       nm = genes)

#get section ids of experiment
retrieve_sections = function(experiment_id) {
  sections <-  read.csv(paste0("http://api.brain-map.org/api/v2/data/query.csv?criteria=model::SectionImage,rma::criteria,[data_set_id$eq", experiment_id, "]"), header = TRUE)
  return(sections$id)
}
#lapply(experiments, function(x) retrieve_sections(x))

for (i in experiments[1:11]) {
  dir.create(paste0("data/ROIMAPer_accuracy/", i, "/"))
}

lapply(experiments[1:11],
       function(x) lapply(retrieve_sections(x), 
                          function(y) download.file(url = paste0(start_url,
                                                                 y,
                                                                 "?downsample=0&view=expression"),
                                                    destfile = paste0("data/ROIMAPer_accuracy/", x, "/", y, "_expression.jpg"),
                                                    mode = "wb"
                          )
       )
)

lapply(experiments[1:11],
       function(x) lapply(retrieve_sections(x), 
                          function(y) download.file(url = paste0(start_url,
                                                                 y,
                                                                 "?downsample=0"),
                                                    destfile = paste0("data/ROIMAPer_accuracy/", x, "/", y, "_ish.jpg"),
                                                    mode = "wb"
                          )
       )
)
```
