## Extended Data 1 for "ROIMAPer: An Open-Source Framework for Rapid and Accurate Atlas-Based Registration of Individual Brain Images in FIJI": in_situ_hybridization_fusing.ijm

```
directory = getDirectory("get the directory");
directory = replace(directory, "\\", "/");
directory_name = File.getNameWithoutExtension(directory);
higher_path = File.getDirectory(directory);

filelist = getFileList(directory);
filelist = Array.sort(filelist);

fused_directory = higher_path + directory_name + "_fused/";
File.makeDirectory(fused_directory);

setBatchMode("hide");
for (i = 0; i < filelist.length; i = i + 2) {
	print("image: " + i + " out of " + filelist.length);
	ish_channel = filelist[i + 1];
	image_name = File.getNameWithoutExtension(directory + ish_channel);
	open(directory + ish_channel);
	run("Split Channels");
	close(ish_channel + " (blue)");
	close(ish_channel + " (green)");
	
	expression_channel = filelist[i];
	open(directory + expression_channel);
	run("Split Channels");
	
	run("Merge Channels...", "c1=[" + ish_channel + " (red)] c2=[" + expression_channel + " (red)] c3=[" + expression_channel + " (green)] c4=[" + expression_channel + " (blue)] create");
	
	rename(ish_channel);
	
	save(fused_directory + image_name + ".tif");
	close(ish_channel);
	run("Collect Garbage");
}
setBatchMode("show");
print("finished");
```
