## Extended Data 1 for "ROIMAPer: An Open-Source Framework for Rapid and Accurate Atlas-Based Registration of Individual Brain Images in FIJI": in_situ_hybridization_measure.ijm

```
directory = getDirectory("get the directory");
directory = replace(directory, "\\", "/");
directory_name = File.getNameWithoutExtension(directory);
higher_path = File.getDirectory(directory);

filelist = getFileList(directory);
filelist = Array.sort(filelist);
run("Set Measurements...", "area mean standard modal min centroid center perimeter bounding fit shape feret's integrated median skewness kurtosis area_fraction stack display redirect=None decimal=9");
setBatchMode("hide");
for (i = 0; i < filelist.length; i++) {
	if (endsWith(filelist[i],".tif")) {
		open(directory + filelist[i]);
		run("To ROI Manager");
		getDimensions(width, height, channels, slices, frames);
		if (channels < 2) {
			run("Split Channels");
			close(filelist[i]);
			run("Merge Channels...", "c1=[" + filelist[i] + " (red)] c2=[" + filelist[i] + " (green)] c3=[" + filelist[i] + " (blue)] create");
			rename(filelist[i]);
		}
		Stack.setChannel(2);
		rois = Array.getSequence(roiManager("count"));
		roiManager("select", rois);
		roiManager("multi-measure append");
		roiManager("reset");
		close(filelist[i]);
	}
}
setBatchMode("show");
selectWindow("Results");
saveAs("results", higher_path + directory_name + "_measured.csv");
close("Results");
print("done");
```
