## Extended Data 1 for "ROIMAPer: An Open-Source Framework for Rapid and Accurate Atlas-Based Registration of Individual Brain Images in FIJI": ROIMAPer_quantification.R

```
library(ggrepel)
library(tibble)
library(GenomicFeatures)
library(AnnotationDbi)
library(org.Mm.eg.db)
library(TxDb.Mmusculus.UCSC.mm10.knownGene)
library(circlize)
library(xml2)
library(scales)
library(XML)
library(tidyverse)
library(ggpmisc)
library(MVN)
library(TOSTER)

files <- list.files("data/ROIMAPer_accuracy/")
regions <- c("Isocortex", "OLF", "HPF", "CTXsp", "STR", "PAL", "TH", "HY", "MB", "P", "MY", "CB")
all_regions <- str_trim(stringr::str_split(readLines(con = "data/ROIMAPer_accuracy/roimapper_accuracy_regions.txt"), pattern = ",")[[1]])
genes = c("pde10a","serpina9","glra1","myl4","tcf7l2","ndst4","gabra6","prox1","mcu", "kcnd2", "sstr4")

experiments = setNames(c(71924204,71670728,72108824,72129251,72339557,74511837,75551467,73520980,72472776,71020432,73636037), 
                       nm = genes)

intensity_list <- list()
for (i in 1:length(genes)) {
  
  filename <- grep(paste0("[0-9]*_", 
                          genes[i], 
                          "_.*combined_measured.csv"), 
                   files, 
                   value = TRUE)
  intensity_list[[i]] <- separate(read.csv(paste0("data/ROIMAPer_accuracy/", 
                                                  filename),
                                           header = TRUE,
                                           row.names = 1), 
                                  "Label", 
                                  sep = ":", 
                                  into = c("Image_Name", "Region", "Channel", "channelname"))
  intensity_list[[i]]$Gene <- genes[i]
}

get_aba_xml <- function(experiment_id) {
  temp <- tempdir()
  temp_file <- paste0(temp, "xml.xml")
  
  url <- paste0("http://api.brain-map.org/api/v2/data/SectionDataSet/query.xml?id=", experiment_id, "&include=structure_unionizes%28structure%29")
  download.file(url, temp_file)
  
  data <- read_xml(temp_file)
  file.remove(temp_file)
  
  structure <- xml_text(xml_find_all(data, ".//acronym"))
  expression_energy <- xml_double(xml_find_all(data, ".//expression-energy"))
  
  return(
    data.frame(
      Region = structure,
      Expression_energy = expression_energy
    )
  )
}

aba_ish_list <- lapply(experiments, function(x) get_aba_xml(x))

for (i in 1:length(genes)) {
  aba_ish_list[[i]]$Gene <- genes[i]
}

#side by side ROIMAPer quantification and Allen Institute
comparison_df <- merge(
  do.call(rbind, intensity_list) %>%
    filter(Region %in% all_regions) %>%
    group_by(Region, Gene) %>%
    summarise(ROIMAPer = mean(Mean)),
  do.call(rbind, aba_ish_list) %>% 
    filter(Region %in% all_regions) %>%
    group_by(Region, Gene) %>%
    summarise(Allen_Institute = mean(Expression_energy)),
  by = c("Region", "Gene")) %>%
  select(Gene, Region, ROIMAPer, Allen_Institute) %>%
  pivot_longer(cols = c("ROIMAPer", "Allen_Institute"), names_to = "Method", values_to = "Signal")

brain_summed_signal <- comparison_df %>%
  filter(Region %in% regions) %>%
  group_by(Method, Gene) %>%
  summarise(Total = sum(Signal))

percentage_df <- merge(
  comparison_df,
  brain_summed_signal,
  by = c("Method", "Gene")) %>%
  mutate(Relative_Signal = Signal / Total) %>%
  pivot_wider(values_from = c("Signal", "Total", "Relative_Signal"), names_from = "Method")

#do Two one-side test of equivalence
tost_list <- list()
for (i in c("myl4", "ndst4", "pde10a", "serpina9", "tcf7l2", "glra1", "gabra6")) {
  x <- percentage_df[percentage_df$Gene == i & percentage_df$Region %in% regions, "Relative_Signal_ROIMAPer"]
  y <- percentage_df[percentage_df$Gene == i & percentage_df$Region %in% regions, "Relative_Signal_Allen_Institute"]
  tost_list[[i]] <- t_TOST(x = unlist(x), 
                           y = unlist(y), 
                           paired = TRUE,
                           hypothesis = "EQU",
                           eqbound_type = "raw",
                           eqb = 0.05,
                           alpha = 0.05/6)
}
tost_list

tost_list_hpf <- list()

for (i in c("sstr4", "mcu", "kcnd2", "prox1")) {
  x <- percentage_df[percentage_df$Gene == i & percentage_df$Region %in% c("CA1", "CA2", "CA3", "DG"), "Relative_Signal_ROIMAPer"]
  y <- percentage_df[percentage_df$Gene == i & percentage_df$Region %in% c("CA1", "CA2", "CA3", "DG"), "Relative_Signal_Allen_Institute"]
  tost_list_hpf[[i]] <- t_TOST(unlist(x), 
                               unlist(y), 
                               paired = TRUE,
                               hypothesis = "EQU",
                               eqbound_type = "raw",
                               eqb = 0.05,
                               alpha = 0.05/3)
}
tost_list_hpf


ish_comparison_plot <- comparison_df %>%
  filter(Region %in% regions, Gene %in% c("myl4", "ndst4", "pde10a", "serpina9", "tcf7l2", "glra1", "gabra6")) %>%
  ggplot(aes(x = factor(Region, levels = regions), 
             y = Signal, 
             fill = Method, 
             group = Method)) +
  geom_col(position = position_dodge2(padding = 0.2)) +
  facet_wrap(vars(factor(Gene, 
                         levels = c("myl4", "ndst4", "pde10a", "serpina9", "tcf7l2", "glra1", "gabra6"))), 
             scales = "free_y", ncol = 1) +
  theme(axis.text.x = element_text(angle = 90, vjust = 0.5, hjust=1),
        legend.position = "bottom",
        legend.title = element_blank(),
        axis.title.x = element_blank()) +
  labs(y = "Mean expression energy intensity")


ish_comparison_plot_hpf <- comparison_df %>%
  filter(Region %in% c("CA1", "CA2", "CA3", "DG"), Gene %in% genes[c(8, 10, 11, 12)]) %>%
  ggplot(aes(x = Region, 
             y = Signal, 
             fill = Method, 
             group = Method)) +
  geom_col(position = position_dodge()) +
  facet_wrap(vars(factor(Gene, levels = c("sstr4", "mcu", "kcnd2", "prox1"))), scales = "free_y", ncol = 1) +
  theme(axis.text.x = element_text(angle = 90, vjust = 0.5, hjust=1),
        legend.title = element_blank(),
        axis.title.x = element_blank()) +
  labs(y = "Mean expression energy intensity")

pdf(file = "figures/in_situ_hybridization_comparison.pdf", width = 3.5, height = 11, fonts = "sans")
ish_comparison_plot
dev.off()

pdf(file = "figures/in_situ_hybridization_comparison_hpf.pdf", width = 4, height = 5.4, fonts = "sans")
ish_comparison_plot_hpf
dev.off()
```
